# A consistent picture of TRPV1 activation emerges from molecular simulations and experiments

**DOI:** 10.1101/310151

**Authors:** Marina A. Kasimova, Aysenur Torun Yazici, Yevgen Yudin, Daniele Granata, Michael L. Klein, Tibor Rohacs, Vincenzo Carnevale

## Abstract

Although the structure of TRPV1 has been experimentally determined in both the closed and open states, very little is known about its activation mechanism. In particular, the conformational changes occurring in the pore domain and resulting in ionic conduction have not been identified yet. Here, we suggest a hypothetical molecular mechanism for TRPV1 activation, which involves the rotation of a conserved asparagine in S6 from the S4-S5 linker toward the pore. This rotation is correlated with the dehydration of four peripheral cavities located between S6 and the S4-S5 linker and the hydration of the pore. In light of our hypothesis, we perform bioinformatics analyses of TRP and other evolutionary related ion channels, analyze newly available structures and re-examine previously reported water accessibility and mutagenesis experiments. Overall, we provide several independent lines of evidence that corroborate our hypothesis. Finally, we show that the proposed molecular mechanism is compatible with the currently existing idea that in TRPV1 the selectivity filter acts as a secondary gate.

## INTRODUCTION

TRPV1 is a member of the transient receptor potential (TRP) family, which promotes non-selective cation current across cell membranes in response to heat, low pH, and inflammatory agents (1–3). Activation of this channel induces burning pain: the action potential triggered by TRPV1 in sensory ganglia and small sensory C-and Aδ fibers is transmitted through the second-order projection neurons in the spinal cord to the thalamus and specific higher order brain areas where it is perceived as pain (4, 5).

TRPV1 is a homotetramer with each subunit being composed of the transmembrane and cytoplasmic regions (6–8). The transmembrane region has six helical segments (S1 through S6); four of them (S1 through S4) constitute a domain structurally homologous to the voltage sensor domain of voltage-gated ion channels. The remaining two helices (S5 and S6) assemble into the pore domain. Cryo-EM structures of TRPV1 in different functional states have revealed key elements of its activation mechanism (6–8). For instance, it became apparent that the gate is located at the crossing of the S6 helical bundle. The gate is closed in the absence of an activating stimulus, while at room temperature it opens upon binding of the agonist capsaicin between S4 and S5 (7). The molecular mechanism of this conformational change has been in part clarified by mutagenesis experiments (8, 9): capsaicin forms a hydrogen bond with the S4-S5 linker and pulls it in the outward direction; as for voltage-gated ion channels, this motion relieves the constriction exerted by the linker on the S6 helical bundle.

However, the cryo-EM structure of the capsaicin-bound state is closed at cryogenic temperatures: the gate is narrower than that of the open state (7), and in simulations it is dehydrated (10). Consistently, Jara-Oseguera et al. have shown that TRPV1 activation becomes increasingly less probable as temperature is decreased even when capsaicin is present in large concentrations (11). Thus, the capsaicin-bound state is closed at cryogenic temperatures and opens as temperature is increased with the mid-point of activation located approximately at 300 K.

We recently used molecular dynamics and free energy calculations to study the molecular mechanism of the closed-to-open transition in the capsaicin-bound state (10). Our major finding was that the conserved asparagine N676 in S6 adopts two alternative conformations in the open and closed states, projecting its side chain towards either the pore or four peripheral cavities located between the S6 helix and the S4-S5 linker, respectively. When TRPV1 is in the closed state, these cavities are connected to the intracellular solution and host several water molecules. Dehydration of the peripheral cavities correlates with the rotation of N676 from these cavities toward the pore of which N676 promotes hydration and thus permeability to ions (Fig. 1).

**Figure 1:**
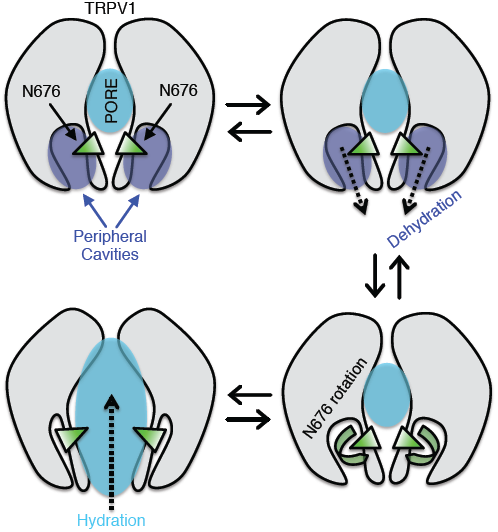
A possible molecular mechanism for TRPV1 activation (10). The channel is shown in grey; water in the pore and peripheral cavities in cyan and blue, respectively; the conserved asparagine N676 in S6 in green. In the capsaicin-bound closed state, the peripheral cavities host several water molecules, while the pore is partially dehydrated; N676 projects its side chain toward the peripheral cavities. Upon application of an activating stimulus, the peripheral cavities dehydrate, an event that correlates with the rotation of N676 toward the pore. The presence of a hydrophilic side chain inside the pore promotes its hydration and hence permeability to ions.

Here, we will briefly summarize our previous computational findings and discuss several independent pieces of experimental evidence that corroborate our hypothesis. We will show how these lend credibility to the rotation of N676 and the presence of the peripheral cavities. Finally, we will present an analysis of our previously reported simulations suggesting that the selectivity filter acts as a secondary gate.

## RESULTS AND DISCUSSION

### The π-bulge in the S6 helix is evolutionarily conserved across the TRP family

We start by recalling a remarkable feature of the capsaicin-bound structure (7): the presence of a π-helical segment between Y671 and N676 (Fig. 2A). α- and π-helices show an important difference: while in α-helices hydrogen bonds join main chain groups that are four residues apart (i-i+4), this spacing changes to five residues in π-helices (i-i+5). Therefore, the presence of a π-helical segment in an α-helix implies a mismatch in the pattern of hydrogen bonds, which results in unpaired carbonyl and amino groups. In practice, one residue does not participate in the network of hydrogen bonds and bulges out of the α-helix forming a so-called π-bulge (Fig. 2A). The π-bulge is mobile: the bulging residue can “reabsorb” by forming hydrogen bonds at the expense of a neighboring residue, which, as a result, bulges out of the helix.

**Figure 2:**
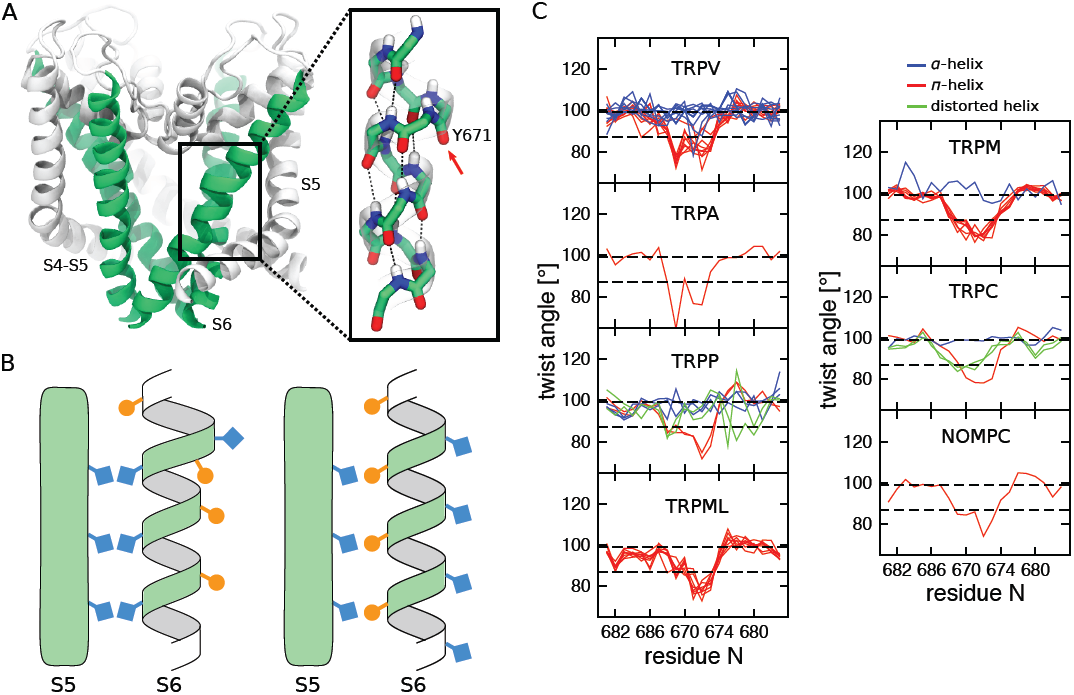
π-helical segment in the S6 helix of TRP channels. A. Cartoon representation of the TRPV1 pore domain (7). The S6 helices are colored in green, and the rest of the pore domain in white. The inset shows the π-helical segment and the π-bulge located on the Y671 residue (highlighted with an arrow). The atoms are colored by element type: C in green, N in blue, O in red, and H in white. B. Schematic representation of an α-helix (right) and an α-helix with a π-bulge (left). Note that the residues facing S5 in the α-helix are lining the pore when the π-bulge is introduced. C. Per residue twist angle in the S6 helices of the available TRP structures estimated using HELANAL (13). The structures with the π-bulge are shown in red (TRPV1, TRPV6, TRPA1, TRPM2, TRPM4, TRPC4, TRPP2, NOMPC, TRPML1 and TRPML3), and without the π-bulge in blue (TRPV2, TRPV4, TRPV5, TRPV6, TRPM8, TRPC3 and TRPP2). The structures in which S6 is distorted and cannot be assigned to either α-helix or α-helix with π-bulge are shown in green (TRPC3, TRPC6 and TRPP2). For the pdb codes of the structures, see Table 1. The twist angle is 99.4° and 86.6° for idealized α-helix and π-helix, respectively (highlighted by dashed black lines).

In one of our previous investigations we hypothesized, on the basis of the sole TRPV1 structure, that this structural motif is conserved across the entire family of TRP channels (12). To test this hypothesis, we analyzed patterns of correlations in a large multiple sequence alignment containing approximately 3000 distinct sequences of genes encoding for TRP channels. The presence of a π-helical segment changes the register of the downstream α-helix: it offsets by one position all the subsequent residues effectively imparting a rotation of ∼100° to the downstream α-helix (Fig. 2B). The change of the register results in a specific pattern of contacts between S6 and the neighboring S5. We detected this pattern in the multiple sequence alignment as strong correlations between S5 and S6 residues; importantly, these pairs of residues are in spatial proximity only with the “correct” register of S6, *i.e.* if the π-bulge is present (12). The conservation of this specific pattern of contacts between S5 and S6 across the entire TRP family suggests that the π-bulge is conserved as well.

**Table 1:** Structures of TRP channels. Columns contain: pdb id, resolution, residue forming the gate, pore radius at the level of the gate, presence/absence of the π-bulge in the S6 helix, conserved asparagine and conformational state of the conserved asparagine (inside or outside the pore). The pore radius at the level of the gate was estimated using HOLE (38). For TRPV2 6bwm and 6bwj structures, the side chain of the residue forming the gate was either not resolved or resolved partially; the pore radius was not estimated for these structures. TBP – to be published.

| TRP channel | PDB ID | Resolution (Å) | Residue forming the gate | R (Å) | π-bulge? | Conserved asparagine | State | Ref |
| --- | --- | --- | --- | --- | --- | --- | --- | --- |
| TRPV1 | 5irz | 2.95 | I679 | 0.7 | yes | N676 | in | (8) |
| TRPV1 | 5is0 | 3.43 | I679 | 0.7 | yes | N676 | in | (8) |
| TRPV1 | 3j5p | 3.275 | I679 | 0.8 | yes | N676 | in | (6) |
| TRPV1 | 3j5r | 4.2 | I679 | 1.8 | yes | N676 | out | (7) |
| TRPV1 | 5irx | 2.95 | I679 | 2.8 | yes | N676 | in | (8) |
| TRPV1 | 3j5q | 3.8 | I679 | 3.0 | yes | N676 | in | (7) |
| TRPV1 | 3j9j | 3.275 | I276 | 1.0 | yes | N273 | in | (14) |
| TRPV2 | 5an8 | 3.8 | M643 | 0.9 | no | N637 | out | (31) |
| TRPV2 | 5hi9 | 4.4 | M645 | 1.1 | no | N639 | out | (32) |
| TRPV2 | 6bwm | 3.9 | M643 not resolved | - | no | N637 | out | (35) |
| TRPV2 | 6bwj | 3.1 | M643 partially resolved | - | no | N637 | out | (35) |
| TRPV4 | 6c8f | 6.5 | M714 | 0.7 | no | N708 | out | (36) |
| TRPV4 | 6c8g | 6.31 | M714 | 0.7 | no | N708 | out | (36) |
| TRPV4 | 6c8h | 6.5 | M714 | 0.7 | no | N708 | out | (36) |
| TRPV4 | 6bbj | 3.8 | M714 | 0.8 | no | N708 | out | (36) |
| TRPV5 | 6b5v | 4.8 | M578 | 0.8 | no | N572 | out | (33) |
| TRPV6 | 6bo8 | 3.6 | I575 | 2.9 | yes | N572 | in | (28) |
| TRPV6 | 6bo9 | 4.0 | I575 | 3.1 | yes | N572 | in | (28) |
| TRPV6 | 6bob | 3.9 | M577 | 0.8 | no | N571 | out | (28) |
| TRPV6 | 6boa | 4.2 | M578 | 0.8 | no | N572 | out | (28) |
| TRPA1 | 3j9p | 4.24 | I957 | 1.4 | yes | N954 | in | (15) |
| TRPM2 | 6co7 | 3.07 | I1082 | 0.9 | yes | N1079 | in | (16) |
| TRPM4 | 6bqv | 3.1 | I1040 | 0.7 | yes | N1037 | in | (17) |
| TRPM4 | 6bqr | 3.2 | I1040 | 0.8 | yes | N1037 | in | (17) |
| TRPM4 | 5wp6 | 3.8 | I1040 | 1.0 | yes | N1037 | in | (18) |
| TRPM4 | 6bcj | 3.14 | I1036 | 0.9 | yes | N1033 | in | (19) |
| TRPM4 | 6bcl | 3.54 | I1036 | 0.8 | yes | N1033 | in | (19) |
| TRPM4 | 6bco | 2.88 | I1036 | 0.9 | yes | N1033 | in | (19) |
| TRPM4 | 6bcq | 3.25 | I1036 | 0.9 | yes | N1033 | in | (19) |
| TRPM8 | 6bpq | 4.1 | L973 | 0.8 | no | N967 | out | (34) |
| TRPC3 | 6cud | 3.3 | I658 | 0.7 | no | N652 | out | (37) |
| TRPC3 | 5zbg | 4.36 | I655 | 1.8 | S6 distorted | N652 | in | (39) |
| TRPC4 | 5z96 | 3.28 | I617 | 0.9 | yes | N614 | in | (20) |
| TRPC6 | 5yx9 | 3.8 | I724 | 1.5 | S6 distorted | N721 | in | (39) |
| TRPP2 | 5mke | 4.3 | L677 | 1.9 | S6 distorted | N674 | in | (40) |
| TRPP2 | 5mkf | 4.2 | L677 | 1.0 | S6 distorted | N674 | out | (40) |
| TRPP2 | 5t4d | 3.0 | L677 | 0.6 | yes | N674 | in | (21) |
| TRPP2 | 5k47 | 4.22 | L677 | 0.7 | yes | N674 | in | (22) |
| TRPP2 | 6d1w | 3.54 | I680 | 2.3 | no | N674 | out | (29) |
| TRPP2 | 5z1w | 3.38 | I560 | 2.6 | no | N554 | out | TBP |
| TRPP2 | 6du8 | 3.11 | I560 | 0.5 | no | N554 | out | (30) |
| NOMPC | 5vkq | 3.55 | I1554 | 0.8 | yes | Q1551 | in | (23) |
| TRPML1 | 5wj5 | 3.72 | I514 | 0.9 | yes | S511 | in | (24) |
| TRPML1 | 5wj9 | 3.49 | I514 | 2.4 | yes | S511 | in | (24) |
| TRPML1 | 5wpq | 3.64 | I514 | 0.9 | yes | S511 | in | (25) |
| TRPML1 | 5wpt | 3.75 | I514 | 0.8 | yes | S511 | in | (25) |
| TRPML1 | 5wpv | 3.59 | I514 | 0.6 | yes | S511 | in | (25) |
| TRPML3 | 6aye | 4.06 | I498 | 0.8 | yes | S495 | in | (26) |
| TRPML3 | 6ayf | 3.62 | I498 | 3.3 | yes | S495 | in | (26) |
| TRPML3 | 5w3s | 2.94 | I498 | 1.0 | yes | S495 | in | (27) |

Since then, several experimental structures of TRP channels have confirmed our hypothesis about the structural conservation of the π-bulge. The structures of TRPV1, TRPV6, TRPA1, TRPM2, TRPM4, TRPC4, TRPP2, TRPN-like channel NOMPC, TRPML1 and TRPML3 all show a π-helical segment in S6 (Fig. 2C and Table 1) (6–8, 14–28). Interestingly, structures without the π-bulge have also been determined for some of these TRP channels (TRPV6 and TRPP2 (28–30)), suggesting that a transition between the π- and α-helical conformations might be part of the functional cycle (Fig. 2C and Table 1). Finally, only TRPV2, TRPV4, TRPV5, TRPM8 and TRPC3 do not show a π-helical segment in S6 (31–37). Either these TRP channels lack altogether the π-helical conformation or, alternatively, this conformation has not been experimentally captured yet.

### The π-bulge causes conformational flexibility of neighboring residues

Analysis of a comprehensive multiple sequence alignment of TRP channels revealed that the pore-lining residues are all hydrophobic except for one that in most of the sequences is an asparagine (N676 in TRPV1, Fig. 3A and B) (12). This feature is strictly conserved in the two evolutionarily divergent families of TRPP and TRPML channels as well. In TRPML channels, instead of an asparagine some sequences have a serine; also in these cases, serine is the only hydrophilic pore-lining residue. The conserved asparagine is located only ∼ 1.5 helical turns away from the π-bulge, which potentially endows this residue with conformational flexibility. Structures of different TRP channels (6–8, 14–37) indeed show that the orientation of N676 (or the corresponding amino acid) with respect to the pore can differ by as much as 180° between two structures (Fig. 3C and Table 1). For instance, in the TRPV sub-family the conserved asparagine points toward the pore in TRPV1 and some structures of TRPV6 (6–8, 14, 28), in the opposite direction (toward the S4-S5 linker) in TRPV2, TRPV4, TRPV5 and the remaining TRPV6 structures (28, 31–33, 35, 36) and, finally, lies just outside the pore in the TRPV1 capsaicin-bound structure (7). Similar conformations are observed in other TRP families including TRPM, TRPC and TRPP (16–22, 29, 30, 34, 37). Importantly, the structures with the conserved asparagine inside the pore or just outside of it all have the π-bulge; by contrast, those in which this residue points to the S4-S5 linker, do not show any π-helical segment in S6 (Table 1).

**Figure 3:**
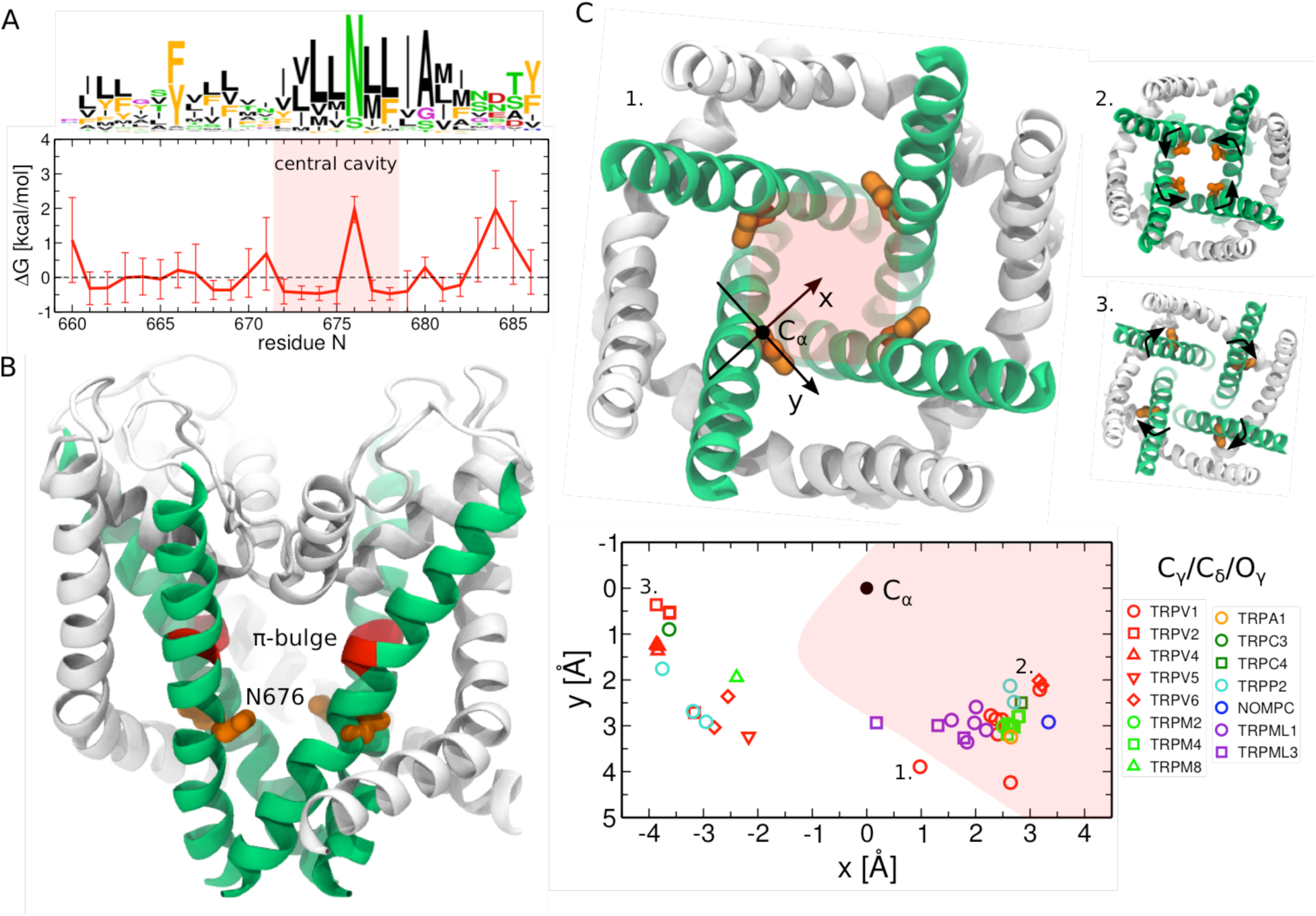
The conserved asparagine adopts a range of conformations from pointing toward the pore to pointing toward the S4-S5 linker. A. Sequence logos and per-position hydrophobicity plot of the S6 helix across the TRP family (numbering based on the TRPV1 sequence) (12). In the sequence logo, the height of each residue at a given position is proportional to its frequency at this position, and the height of the overall stack of residues decreases linearly with Shannon entropy (41). Note that in few cases the conserved asparagine can be substituted by a serine (TRPML channels) or glutamine (TRPN-like channel NOMPC). In the hydrophobicity plot, positions with positive Δ*G* are hydrophilic, and those with negative Δ*G* hydrophobic. The red shaded area highlights the residues lining the central cavity (between Y671 and I679). B. Cartoon representation of the TRPV1 pore domain (7). The S6 helices are shown in green, and the rest of the pore domain in white. The π-bulge and the neighboring conserved asparagine are colored in red and orange, respectively. C. Orientation of the conserved asparagine with respect to the pore in different TRP structures (6–8, 14–37). The side chain position in the plane perpendicular to the pore axis is shown: the Cα-atom is represented as a black dot and is located at (0,0), and the terminal Cγ-atom (Cδ and Oγ in NOMPC and TRPML channels, respectively) is shown as a colored symbol (a different one for each TRP channel). X-axis is aligned with the vector connecting the Cα-atom and the center of the pore. The red shaded area highlights the pore region. The three insets show the pore domains of the TRPV1 capsaicin-bound structure (1), TRPV6 (2) and TRPV2 (3): the conserved asparagine points toward the S4-S5 linker in TRPV2, toward the center of the pore in TRPV6 and lies just outside the pore in the TRPV1 capsaicin-bound structure. In TRPV6 and TRPV2 (the two extreme cases), the black arrows highlight the direction of the asparagine rotation between the conformations inside and outside the pore.

This structural heterogeneity is per se striking: evolutionary conservation points to functional relevance, yet the residue lacks a well-defined conformation. Is the role of the conserved asparagine related to its flexibility? In our previous study (10), we have shown that in the TRPV1 capsaicin-bound structure the rotation of N676 inside or outside the pore significantly affects the polarity of the latter and hence ionic transport: only when N676 is exposed to the pore, the latter is continuously hydrated and the free energy barrier for ionic transport at the level of the gate is ∼ 5.2 kcal/mol. By contrast, when N676 is interacting with the S4-S5 linker, the free energy barrier is larger than 12 kcal/mol, thereby preventing ionic transport. Consistent with these predictions, two recently determined experimental structures of TRPV6 show that N572 faces the pore in the open conformations and the S4-S5 linker in the closed one (28).

It is important to note that the location of the conserved asparagine side chain inside the pore may not be the only determinant of channel opening. The radius of the pore should be, in any case, large enough to permit the passage of hydrated ions. Accordingly, several structures of TRP channels showing the conserved asparagine in the pore-facing configuration and the pore radius smaller than 2 Å are likely representatives of the closed state (Table 1).

### Y671 motion affects gating at the selectivity filter and is coupled to N676 rotation

Is there functional data showing that the rotation of S6 underlies TRP channels’ gating? Steinberg et al. recently reported that TRPV1 activation correlates with a conformational change at the level of the π-bulge where a non-natural amino acid bearing a coumarin side chain was introduced (42). Coumarin is a fluorophore with exquisite sensitivity to environmental polarity (43, 44); when genetically encoded into proteins, it can be used as reporter of changes in local hydration and conformational rearrangements. In the TRPV1 channel with Y671 substituted for coumarin, activation by capsaicin was found to cause an increase in both the photon counts and optical fluctuations. This effect suggests that coumarin undergoes a conformational transition from a more hydrated state to a less hydrated one (42).

While the exact conformational change cannot be inferred by these experiments, molecular dynamics simulations performed on the TRPV1 wild type suggested a possible explanation for the observed changes in fluorescence (42). The simulations showed that the surface area of Y671 accessible to solvent decreases from 30% to 15% on passing from the closed to open states, in agreement with the experimental data for coumarin. The reason for this change is the conformational rearrangement of Y671 coupled to gating through N676. In the TRPV1 open state, Y671 is oriented perpendicularly with respect to the pore axis and it is shielded from water molecules by the pore helices. The N676 side chain is located inside the pore where it forms a hydrogen bond with the I672 carbonyl group, which stabilizes the π-bulge on I672 (Fig. 4A). In the closed state, the N676 side chain moves toward the S4-S5 linker and no longer donates a hydrogen bond to the main chain. As a consequence, the π-bulge is found preferentially on Y671 (Fig. 4A). The displacement of the π-bulge from I672 to Y671 causes a change of Y671 orientation with respect to the pore: it goes from being perpendicular to the pore axis in the open state to being parallel to it in the closed state (Fig. 4A and B). In the conformation parallel to the pore axis, Y671 becomes less shielded by the pore helices and hence more accessible to water molecules (Fig. 4B).

**Figure 4:**
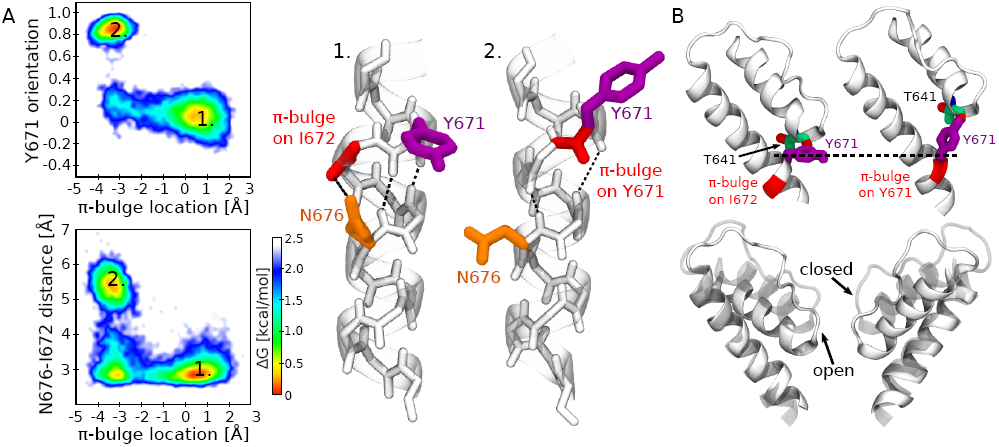
Two different conformations of the S6 helix in the TRPV1 closed and open states. A. Coupling between N676 and Y671. The top left panel shows Y671 orientation (the cosine of the angle between the Cα-Cγ vector and the pore axis) and the π-bulge position (the difference between the two distances: I672 carbonyl oxygen – N676 amine hydrogen and Y671 carbonyl oxygen – N676 amine hydrogen; positive values indicate the location of the π-bulge on I672, while negative ones indicate that the π-bulge is on Y671). Only two conformations are observed: open state (1), Y671 is perpendicular to the pore axis, and the π-bulge is on I672; closed state (2), Y671 is parallel to the pore axis, and the π-bulge is on Y671. The bottom left panel shows the distance between N676 carboxamide carbon and I672 carbonyl oxygen and the π-bulge position. The energetically most favorable conformations are: open state (1), a hydrogen bond between N676 carboxamide and I672 carbonyl oxygen is present, and the π-bulge is on I672; closed state (2), the hydrogen bond between N676 carboxamide and I672 carbonyl oxygen is absent, and the π-bulge is on Y671. The right panel shows the cartoon representations of the two conformations. Y672 is colored in purple, the π-bulge in red, and N676 in orange. The dashed lines show the network of hydrogen bonds. B. A change of Y671 orientation results in a displacement of the pore helix. The top panel shows the conformations of Y671 and of the adjacent pore helix in the open (left) and closed (right) states. The bottom panel shows the superposition between the pore helix in the open (solid) and closed (transparent) states.

Interestingly, the molecular dynamics simulations also suggest that the observed re-orientation of Y671 is coupled to the conformational change of the selectivity filter. In particular, when Y671 side chain is parallel to the pore axis, it donates a hydrogen bond to the T641 carbonyl group on the pore helix. This interaction displaces the pore helix toward the extracellular solution resulting in a slight constriction of the selectivity filter. In our previous study we have shown that this constriction is sufficient to increase the free energy barrier for ionic transport from ∼ 1.5 in the open state to ∼ 5.0 kcal/mol in the closed one (10). Thus our findings suggest that in TRPV1 the selectivity filter can act as a gate whose opening is coupled to the conformation of Y671 and N676. Can this coupling mechanism be conserved in other TRP channels? The analysis of the multiple sequence alignment argues against this hypothesis by showing a lack of conservation at position 671 (Fig. 3A). Moreover, recent structural data indicates that in TRPV4 and TRPML3, the selectivity filter adopts the same conformation in the closed and open states (26, 27, 36).

### Are the peripheral cavities the missing piece of the puzzle?

The molecular mechanism suggested here agrees with the one emerging from the TRPV6 structures (28) and complements it with a detailed description of the conformational rearrangements involved in activation: the rotation of the conserved asparagine increases the hydrophilicity of the pore surface thereby promoting the hydration of this compartment. An interesting finding provided by the molecular dynamics simulations and not anticipated by these structures is the factor stabilizing the closed state, in which the asparagine is located outside the pore. Our simulations have shown that in the closed state of TRPV1 N676 is accommodated inside a small hydrophobic cavity located between the S4-S5 linker and S6 and connected to the intracellular solution (10). This cavity, which we call peripheral in contrast to the central cavity along the ionic pathway, hosts several water molecules forming hydrogen bonds with N676 carboxamide. Importantly, the hydration of the peripheral cavity and the N676 rotation are correlated (10): when the peripheral cavity dehydrates, N676 rotates inside the pore. Based on this observation, we suggested that at least for some stimuli activation of TRPV1 is triggered by the dewetting of the peripheral cavities (10). Is there any experimental evidence that supports the existence of these peripheral cavities? Overall, we found three independent observations, which can be retrospectively interpreted in light of this suggested molecular mechanism: (i) the experiments probing S6 accessibility to intracellular solution (45), (ii) the site-directed mutagenesis experiments targeting S6 residues (46), and (iii) the structural studies of the evolutionary related voltage-gated sodium and calcium channels (47–60).

In a milestone paper by Salazar et al. (45), the authors probed the accessibility of the TRPV1 pore residues to the intracellular solution; they mutated, one by one, all the S6 residues to cysteine and assessed their accessibility to thiol-modifying reagents in the open and closed states. Their results showed that modification of D654-Y671 is more prominent in the open state than in the closed one, and that of M682-G683 is state-independent. Interestingly, the rest of S6, namely the I672-L681 segment, showed different accessibility only when a bulky reagent was used; application of small reagents in the open or closed states in contrast resulted in a similar effect. Based on this data, the authors suggested the presence of lower and upper gates located at the level of L681 and Y671, respectively; both gates are open in the open state, while in the closed state the upper gate is closed and the lower gate is partially open (45). This elegant explanation, however, turned out to be inconsistent with the structural information subsequently obtained via cryo-EM on the apo state (6), in which the gate at the level of the S6 bundle crossing was found to be tightly constricted with a radius of ∼ 1.0 Å, *i.e.* impermeable to any solute.

Can the presence of the peripheral cavities solve this apparent contradiction? To answer this question, we characterized the accessibility of the TRPV1 pore residues in the open and two closed states, including capsaicin-bound closed and apo closed (Fig. 5). In agreement with Salazar et al., we found that all of the S6 residues are accessible to the intracellular solution in the capsaicin-bound open state. Moreover, we found that the same set of residues, namely the I672-G683 segment, is accessible in the capsaicin-bound closed state. Despite the constriction at the level of I679, the residues up to I672 (including) do interact with the intracellular solution. However, the interacting water molecules are not located in the pore, but rather in the peripheral cavities. Therefore, considering that the pore residues can be accessible to the intracellular solution through the peripheral cavities, one can resolve the contradiction between the water accessibility experiments (45) and the structural information (6).

**Figure 5:**
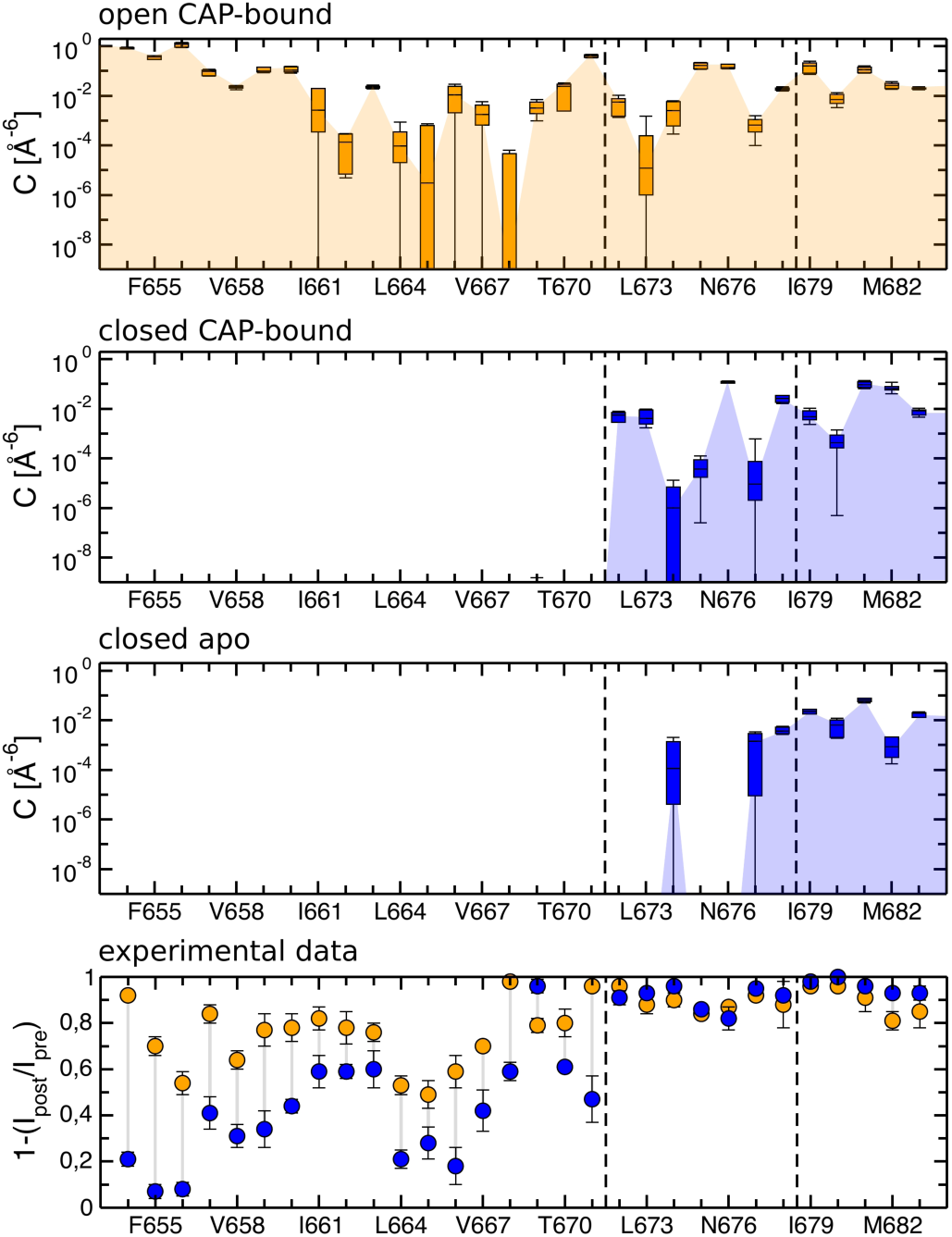
Accessibility of the S6 residues to the intracellular solution in the TRPV1 open (orange) and closed (blue) states. The three top panels show the computational results, while the bottom panel shows the experimental data extracted and digitalized from (45). In the simulations, water accessibility is calculated as the overlap between the atomic density of a given S6 residue and that of the intracellular water (the values shown correspond to a sum over all four channel subunits). The open state data corresponds to the results obtained for the capsaicin-bound open state (10); the closed state data corresponds to the results obtained for the capsaicin-bound closed state and the apo closed state (6, 10). The box plots report median and interquartile range with the whiskers representing the standard deviation. The threshold for water accessibility was chosen to be zero. In the experiments, accessibility to the intracellular solution is measured as an effect on current after application of a thiol-modifying reagent. The dashed lines highlight the hypothetical upper gate according to Salazar et al. (45) and I679 (the gate at the level of the S6 bundle crossing).

Interestingly, several pore residues above the gate are also accessible in the apo closed state, and similar to the capsaicin-bound closed state they interact with the water molecules present in the peripheral cavities. However, in the apo state, the peripheral cavities are significantly smaller and host fewer water molecules. We hypothesize that this difference is the result of the S4-S5 linker movement: upon binding capsaicin pulls the S4-S5 linker in the outward direction thus expanding the space between the S4-S5 linker and S6 (8, 9). The smaller size of the peripheral cavities in the apo closed state explains why only some of the I672-G683 residues are accessible to the intracellular solution.

Another piece of evidence lending credibility to our hypothesis comes from the alanine-scanning mutagenesis of the S6 residues performed by Susankova et al. (46). Susankova et al. have identified a clear periodic pattern of the functional effects of mutations, which is consistent with the α-helical structure of S6 (46). Subsequently obtained cryo-EM structures of TRPV1 showed that the residues whose mutation has the most significant effect on activation are located outside of the pore and line the surface of the peripheral cavities (Fig. 6). Importantly, the response of the corresponding mutants to capsaicin, voltage and heat was significantly reduced, indicating stabilization of the closed state versus the open one. A possible interpretation for this data is that the substitution of hydrophobic residues (I672, L673, L674, L678 and M682) with a less hydrophobic one (alanine) promotes the hydration of the peripheral cavities and thus stabilizes the closed state.

**Figure 6:**
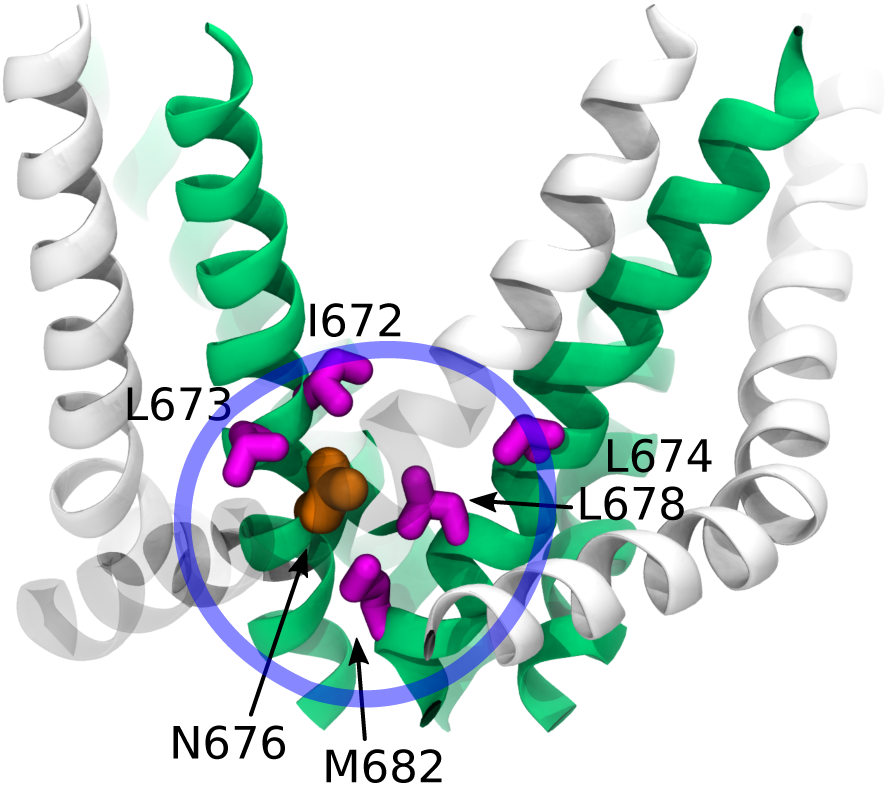
Mutagenesis of S6 residues lining the peripheral cavities. The residues mutated by Susankova et al. are shown in magenta (46), the conserved asparagine in orange, the S6 helices and the rest of the pore domain in green and white, respectively. The peripheral cavity is highlighted with a blue circle.

We finally analyzed the families of voltage-gated sodium and calcium channels (Nav and Cav) that are structurally homologous and evolutionary related to TRP channels (47–60). Interestingly, these families also show a predominantly hydrophobic pore with the only exception of a conserved asparagine at the position corresponding to N676 (Fig. 7A). In NaChBac, mutagenesis of this residue into several other amino acids yielded functional channels only in the case of N225D mutant; and even in that case gating of N225D was drastically affected (61). Furthermore, the structures of Nav and Cav channels show that the conserved asparagine can adopt both the pore inward-facing (NavPas – all subunits, NavEel – one subunit out of four) and outward-facing conformations (Cav1.1 – all subunits, NavAb, NavRh, NavMs, NavEel – three subunits out of four) (Fig. 7B and C) (47–60). Do these channels have the peripheral cavities? A recent high-resolution structure of the bacterial sodium channel NavMs shows the presence of four cavities, whose location coincide with that of the peripheral ones in TRPV1 (56). Moreover, similar to the peripheral cavities, they host the side chain of the conserved asparagine and, importantly, few water molecules interacting with it.

**Figure 7:**
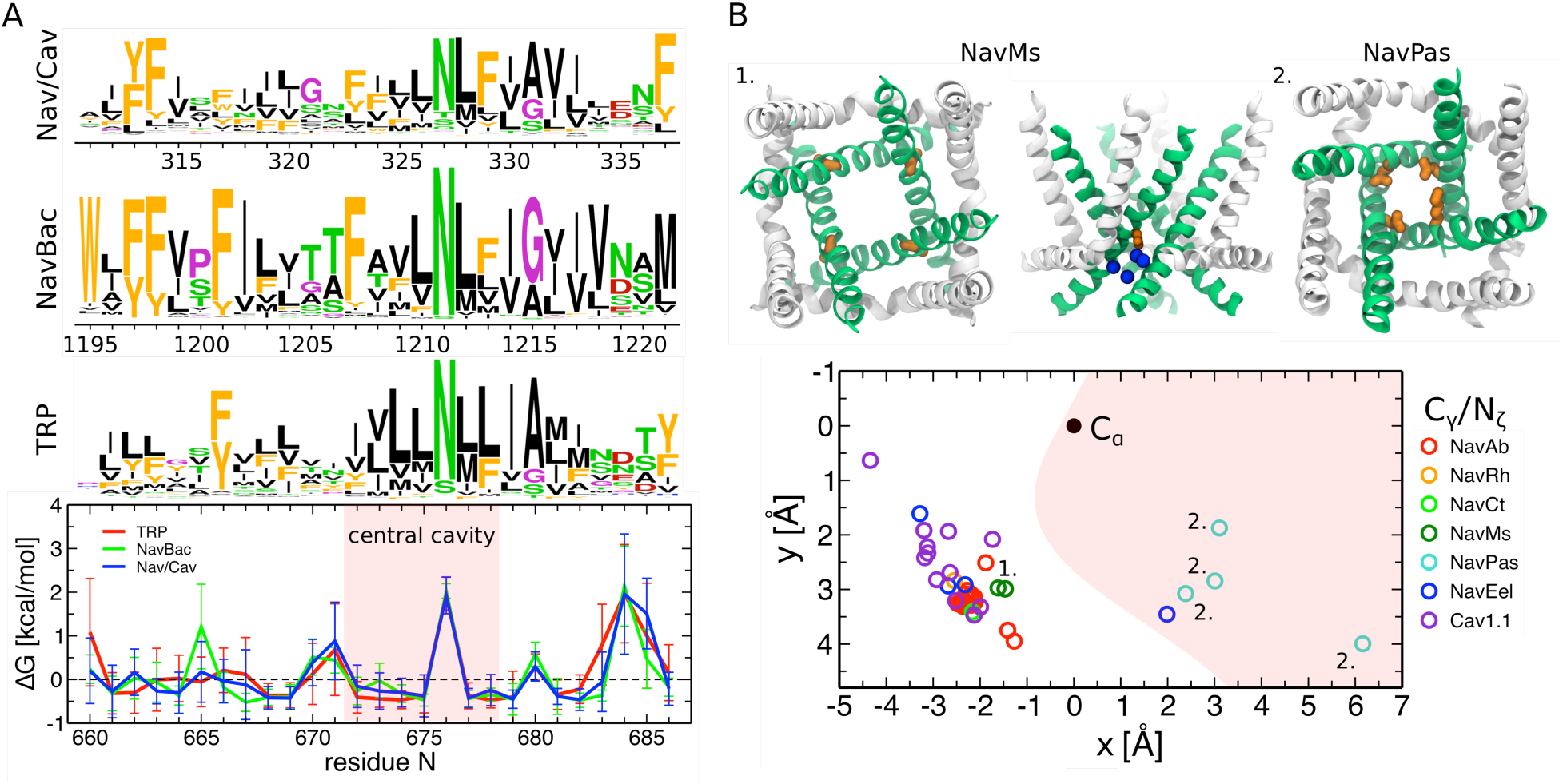
Structurally homologous and evolutionary related families of voltage-gated sodium and calcium channels show a conserved asparagine in S6 and the presence of four hydrated cavities between S6 and the S4-S5 linker. A. Sequence logos and per-position hydrophobicity plots of the S6 helix for voltage-gated sodium and calcium channels (Nav/Cav), bacterial voltage-gated sodium channels (NavBac), and TRP channels (TRP) (12, 62). In the sequence logos, the height of each residue at a given position is proportional to its frequency at this position, and the height of the overall stack of residues is decreases linearly with Shannon entropy (41). In the hydrophobicity plots, positions with positive Δ*G* are hydrophilic, and those with negative Δ*G* hydrophobic. The red shaded area highlights the residues lining the central cavity (between Y671 and I679). B. Orientation of the conserved asparagine with respect to the pore in different Nav and Cav structures (47–60). The side chain position in the plane perpendicular to the pore axis is shown: the Cα-atom is represented as a black dot and is located at (0,0), and the terminal Cγ-atom (Nζ in one of the NavPas subunits where the conserved asparagine is substituted for a lysine) is shown as colored symbols (different for each Nav/Cav channel). X-axis is aligned with the vector connecting the Cα-atom and the center of the pore. The red shaded area highlights the pore region. The two insets show the pore domains of the NavMs structure (1, top and side views) and of the NavPas structure (2, top view). The conserved asparagine points toward the S4-S5 linker in NavMs and toward the pore in NavPas. In NavMs, the water molecules inside the cavity located between the S6 helix and the S4-S5 linker are shown in blue.

## CONCLUSIONS

Despite two decades of extensive investigations, the molecular mechanism of TRPV1 activation is only partially understood. Catching an ion channel in the act of gating is, in general, a complex task requiring techniques with both high temporal and high spatial resolution. NMR spectroscopy and fluorescence/luminescence based approaches are typically the tools of choice but they cannot easily probe the motions of an ion channel when embedded in a cell membrane. This limitation is particularly severe in the case of TRPV1 whose activity shows exquisite dependency on numerous environmental factors, such as lipid composition (8, 63–67), extracellular concentration of Na^+^ (11) and the presence of cholesterol or other fatty acids (68–71). Accordingly, advanced experimental techniques, such as cell unroofing (72, 73) and incorporation of novel unnatural amino acids (42, 73), are being constantly developed to overcome these limitations and introduce only minimal perturbations to the native environment of the channel. Despite the great progress, these techniques have focused so far only on specific aspects of TRPV1 activation. A global picture of this molecular process is still lacking and it is unlikely that a single experimental technique will provide it in the foreseeable future.

Here, prompted by our previous molecular dynamics simulations (10) and bioinformatics analyses (12), we propose a molecular mechanism for TRPV1 activation. More than definite answers, our contribution provides testable predictions and an overall framework to rationalize a wealth of existing experimental data. Indeed, despite being consistent with the results of several independent studies, our hypothesis is still in part speculative. Most importantly, our work was only focused on the capsaicin-bound state, and the transition between that and the apo one was not investigated. It remains unclear how the conformation of the peripheral cavities changes along this transition and how capsaicin binding triggers this change. Based on previous studies (8, 9), we surmise that capsaicin promotes the expansion of the cavities between S6 and the S4-S5 linker. Enhanced sampling simulations starting from the apo state could provide answers to this question and also assess possible temperature dependency of the apo-to-capsaicin-bound transition.

Further research will be required to validate the molecular mechanism proposed here. Discriminative experiments excluding multiple alternative interpretations are highly desirable. For instance, incorporating unnatural amino acids in TRPV1 could allow for a fine-tuning of the polar properties of the pore and the peripheral cavities without perturbing too much their geometry. The same approach could be used to investigate the coupling between the selectivity filter and the gate. According to our model, this coupling involves the interaction between the side chain of N676 and the S6 helix main chain. The latter could be modified by introducing appropriately designed unnatural amino acids. From the computational point of view, long unbiased simulations could be used to sample gating events and provide direct confirmation of the activation mechanism. The Anton 2 supercomputer developed by D. E. Shaw Research (74, 75) and available through the Pittsburgh Supercomputing Center provides the opportunity to reach timescales comparable to those required for TRPV1 activation and thus to test the hypothesized chain of events. These simulations ought to involve, ideally, both the apo and capsaicin-bound states.

In closing, it is interesting to comment on an intriguing possibility emerging from our analyses: we found that the key structural features supporting our hypothesized mechanism are present also in the evolutionary related families of voltage-gated sodium and calcium channels. In particular, the conserved asparagine in the S6 helix, the π-bulge and the peripheral cavities are seemingly conserved beyond the family of TRP channels. This raises the possibility that voltage-gated sodium and calcium channels might share with TRP channels important aspects of the activation mechanism.

However, the sequence and structure conservation of S6 across TRP and voltage-gated sodium and calcium channels highlights a potentially problematic aspect of our hypothesis: in our previous computational investigation (10), we suggested that the rotation of N676 underlies temperature sensitivity in TRPV1. The evolutionary conservation of this asparagine cannot be easily reconciled with the fact that several TRP and voltage-gated sodium and calcium channels are insensitive to temperature. Future investigations ought to improve our understanding of temperature sensitivity of ion channels and thus contribute to solve this conundrum.

## MATERIALS AND METHODS

### Molecular dynamics simulations

The molecular dynamics trajectories of the capsaicin-bound closed and open states were taken from our previous study (10). Briefly, the structure of the TRPV1 capsaicin-bound state (7) with four capsaicin molecules (76) was embedded in a hydrated 1-palmitoyl-2-oleoylphosphatidylcholine, POPC, bilayer and surrounded by 150 mM NaCl solution. From 4 to 6 water molecules were added inside each peripheral cavity in one case (this system further relaxed to the closed state), while these cavities were left empty in the other case (this system further relaxed to the open state) (10). The CHARMM36 force field (77) was used for the protein and the POPC lipids, the parameters derived in (76) were used for capsaicin, and the TIP3P model (78) was used for water molecules. The molecular dynamics simulations were performed following a multistep protocol (for the details, see (10)) using NAMD 2.10 (79). The pressure and temperature were kept constant at 1 atm and 300 K, respectively, using Langevin dynamics. A cutoff of 11 Å with a switching function between 8 and 11 Å was used for the vdW interactions. A cutoff of 11 Å was considered for the short-range component of electrostatic interactions, and the long-range component was computed using Particle Mesh Ewald (80). The overall length of the molecular dynamics trajectory for each state was 750 ns. In addition, we performed the molecular dynamics simulations of the TRPV1 apo state (54) using an analogous setup. The overall length of the molecular dynamics trajectory for this state was 200 ns.

### Analysis of sequence conservation

The multiple sequence alignments of the TRP and voltage-gated sodium and calcium channels were taken from our previous studies (12, 62). To analyze the sequence conservation for these families we computed the Shannon entropy profile using the logos website (41).

### Estimation of the accessibility of S6 residues to the intracellular solution

To estimate the accessibility of S6 residues to the intracellular solution, we considered the second halves of the molecular dynamics trajectories. We separated the intracellular solution from the rest applying the following procedure. We first calculated the three-dimensional map of water occupancy using the Volmap tool of VMD (81). Based on this we generated a binary map by setting each voxel to either 0 or 1 depending on whether or not the local value of water occupancy overcomes a predefined threshold. After that we used gmx cluster of Gromacs 5.0.4 (82) to group the voxels. This allowed us to identify the cluster containing the intracellular solution and all the regions connected to it. Finally, as a measure of accessibility, we calculated the overlap between the occupancy of the intracellular solution and that of each S6 residue.

## ACKNOWLEDGMENTS

This work was partially supported by National Institutes of Health through grants R01GM093290, S10OD020095, NS055159 and P01GM055876 and National Science Foundation through grants ACI-1614804 and CNS-1625061.

